# *In Utero* MRI of Radial Glial Fibers Highlights Their Role in Modulating Fetal Cortical Folding

**DOI:** 10.64898/2026.10.28.761462

**Authors:** Ruike Chen, Xinyi Xu, Qinfeng Zhu, Weizeng Zheng, Yao Shen, Ziyang Yu, Mingyang Li, Zhiyong Zhao, Cong Sun, Keqing Zhu, Guangbin Wang, Qiong Luo, Dan Wu

**Affiliations:** Key Laboratory for Biomedical Engineering of Ministry of Education, Department of Biomedical Engineering, College of Biomedical Engineering & Instrument Science, Zhejiang University, Hangzhou 310027, China; Department of Radiology, Women’s Hospital School of Medicine Zhejiang University, Hangzhou 310006, China; Children’s Hospital, Zhejiang University School of Medicine, National Clinical Research Center for Child Health, Zhejiang University, Hangzhou, 310003, China; Department of Radiology, Beijing Hospital, National Center of Gerontology, Institute of Geriatric Medicine, Chinese Academy of Medical Sciences, Beijing, 100730, China; National Health and Disease Human Brain Tissue Resource Center, Zhejiang University, Hangzhou 310058, China; Department of Radiology, Shandong Provincial Hospital Affiliated to Shandong First Medical University, Jinan 250021, China; Department of Obstetrics, Women’s Hospital School of Medicine Zhejiang University, Hangzhou 310006, China

## Abstract

The process of cortical folding is fundamental to early human brain development. While the differential growth of the cortical layers is a known driver, the biomechanical contribution of key cellular microstructures remains unclear, and there is a lack of direct *in vivo* markers of cortical microstructures for time-resolved observations. Using a multimodal approach combining *in utero* and *ex vivo* fetal brain MRI, histology, and computational mechanical simulation, we investigated whether developmental reorganization of cortical radial microstructures, particularly the radial glial cell (RGC)-associated scaffold, is linked to cortical folding and may modulate its mechanical progression. We characterized this radial organization noninvasively using diffusion MRI-based radial fiber density (*FD*_rad_). *Ex vivo* fetal brain MRI and histology supported that regional variations in *FD*_rad_ were strongly associated with vimentin-labelled radial architecture. Our *in utero* MRI data revealed closely coordinated developmental changes in cortical *FD*_rad_ and curvature, with their regional timings and spatial patterns significantly coupled across the cortex. Longitudinal *in utero* MRI data further supported the synchronized within-individual co-development of *FD*_rad_ and curvature. Furthermore, biomechanical simulations showed that reducing radial reinforcement modulated differential-growth-driven buckling, producing earlier folding and a greater cortical curvature under otherwise matched model conditions. Collectively, this study introduces *FD*_rad_ as a noninvasive *in utero* MRI measure sensitive to radial glial fibers. By integrating spatial, temporal, histological, and computational evidence, our findings suggested a model in which developmental disassembly of the radial scaffold may modulate the mechanical response of the fetal cortex to differential growth, thereby contributing to cortical folding.

## 1 Introduction

Cortical folding, the process by which the cerebral cortex develops convoluted gyri and sulci, is one of the most prominent morphological events during fetal brain development. Understanding its underlying mechanisms has long been a major challenge in neuroscience. Most existing models are grounded in the “differential growth” theory, positing that the faster expansion of the cortical plate (CP) relative to subcortical tissue drives surface buckling^1,2^. Other studies have incorporated subcortical fiber development to further examine the role of fibers in cortical morphogenesis^3–7^. On the other hand, the contribution of microstructural reorganization within the cortex itself, e.g., developmental changes in glial, dendritic, and neuronal organizations, may also contribute substantially to the folding process.

Histological and genetic studies have provided some of the most direct insights. For example, Wang, et al.^8^ reported an association between dendritic arborization and cortical curvature change, while del Toro, et al.^9^ suggested that the regulation of intracellular adhesion of radially migrating neurons is critical in sulcation. Despite their value, existing studies have rarely looked into the role of cortical radial glial fiber reorganization in cortical folding. Moreover, the histological approach is incapable of capturing time-resolved developmental trajectories in the human brain. Longitudinal *in vivo* observations are desired to understand the highly dynamic folding process. Magnetic resonance imaging (MRI) offers a powerful, noninvasive tool to investigate brain development *in vivo* and longitudinally^10^. Particularly, diffusion MRI (dMRI) enables probing of tissue microstructures by measuring restricted and anisotropic water diffusion in the brain^11^.

Both *in utero* fetal and neonatal dMRI have revealed a key tissue microstructural transformation accompanying the onset of gyrification, i.e., the gradual loss of cortical radial organization during cortical folding, reflected by reductions in fractional anisotropy (FA)^12,13^. These neuroimaging patterns arise from the reorganization of radial glial fibers and the dissolution of columnar neuronal organization as neuronal migration nears completion^14^. Notably, recent evidence suggested that these changes precede the initiation of folding itself^15^. In atypical development, including preterm birth^16^, polymicrogyria^17^, lissencephaly^18^, and autism spectrum disorder^19,20^, MRI studies have further demonstrated that aberrant gyrification patterns are accompanied by concurrent abnormalities in fiber organization. Our previous study linking MRI with regional gene expression suggested that genes up-regulated in regions of higher cortical curvature highlighted SYNE2, which is significantly enriched in the biological process of neuronal migration along radial glial cell (RGC)-associated architecture^21^. These findings suggested a close relation between the RGC-related fiber processes and cortical morphogenesis. Still, the mechanism by which glial fibers modify cortical gyrification remains unclear.

During fetal brain development, RGCs project long radial fibers that span the cortical wall. Collectively, these densely packed fibers constitute a macroscopic radial scaffold that physically guides neuronal migration and mechanically sustains cortical morphology^22,23^. Although individual glial cells may be softer than their neighboring neurons^24^, the collective presence of RGCs contributes significantly to overall tissue stiffness^25^. This is likely attributable to their role in assembling the surrounding extracellular matrix (ECM) and radially organizing neurons^26^, as well as their abundance of vimentin and the tension-induced stiffening maintained while attached to the cerebral wall^27,28^. Upon the completion of neuronal migration, RGCs lost their attachment to the pial and ventricular surfaces and transformed into immature astrocytes^29^, possibly making the cortical tissue more compliant^30,31^. An animal model also observed a transient cortical stiffness decrease in the embryonic brain^25^.

Informed by the existing evidence, we hypothesized that developmental changes in RGC-associated radial organization are coupled with cortical folding and may modulate the cortical mechanical response to differential growth. In this study, we combined *in utero* dMRI, immunohistology, and mechanical simulation to systematically explore the relationship between radially organized cortical microstructures and cortical folding. First, *ex vivo* MRI and histological experiments showed that the dMRI-based radial fiber density (*FD*_rad_) was strongly associated with regional variations in vimentin-labelled radial architecture, supporting its sensitivity to radial glial fiber organization. Using *FD*_rad_ in our *in utero* dMRI dataset (ZJU-Fetal), we observed spatially and temporally coordinated changes between cortical radial fiber organization and cortical folding during mid-gestation. Longitudinal *in utero* MRI from the developing Human Connectome Project (dHCP) was then utilized to further examine this relationship in an independent dataset, revealing coupled within-interval changes in radial organization and cortical curvature. Finally, we used the *in utero* measured cortical microstructural properties to inform computational mechanical simulations, testing the mechanical plausibility that reduced radial reinforcement could modulate differential-growth-driven cortical folding under the assumed constitutive framework. This multi-modal approach provides new insights into how cortical microstructure—specifically, the developmental reduction in radial organization associated with RGC fibers—may contribute to the evolving biomechanical context of cortical folding.

## 2 Results

### 2.1 Histological Support for dMRI-based *FD*_rad_ as a Marker of RGC

#### Fibers

This study evaluated dMRI-based *FD*_rad_, defined as apparent fiber density weighted by the alignment of fiber orientations with the cortical surface normal, as a candidate imaging measure sensitive to radial fiber organization in the fetal cortex. To examine the histological basis of *FD*_rad_, we assessed its regional association with immunohistological measures in an *ex vivo* fetal brain sample. After *ex vivo* ultra-high-resolution dMRI of the 23-week fetal brain sample (Fig. 2d), the brain was sectioned for immunohistology using hematoxylin as a nuclear counterstain, vimentin to label RGC-associated structures, Neurofilament to label axons, and MAP2 to label neuronal dendrites and relatively mature neuronal soma in the deep layers of CP^32^.

**Fig. 1.**
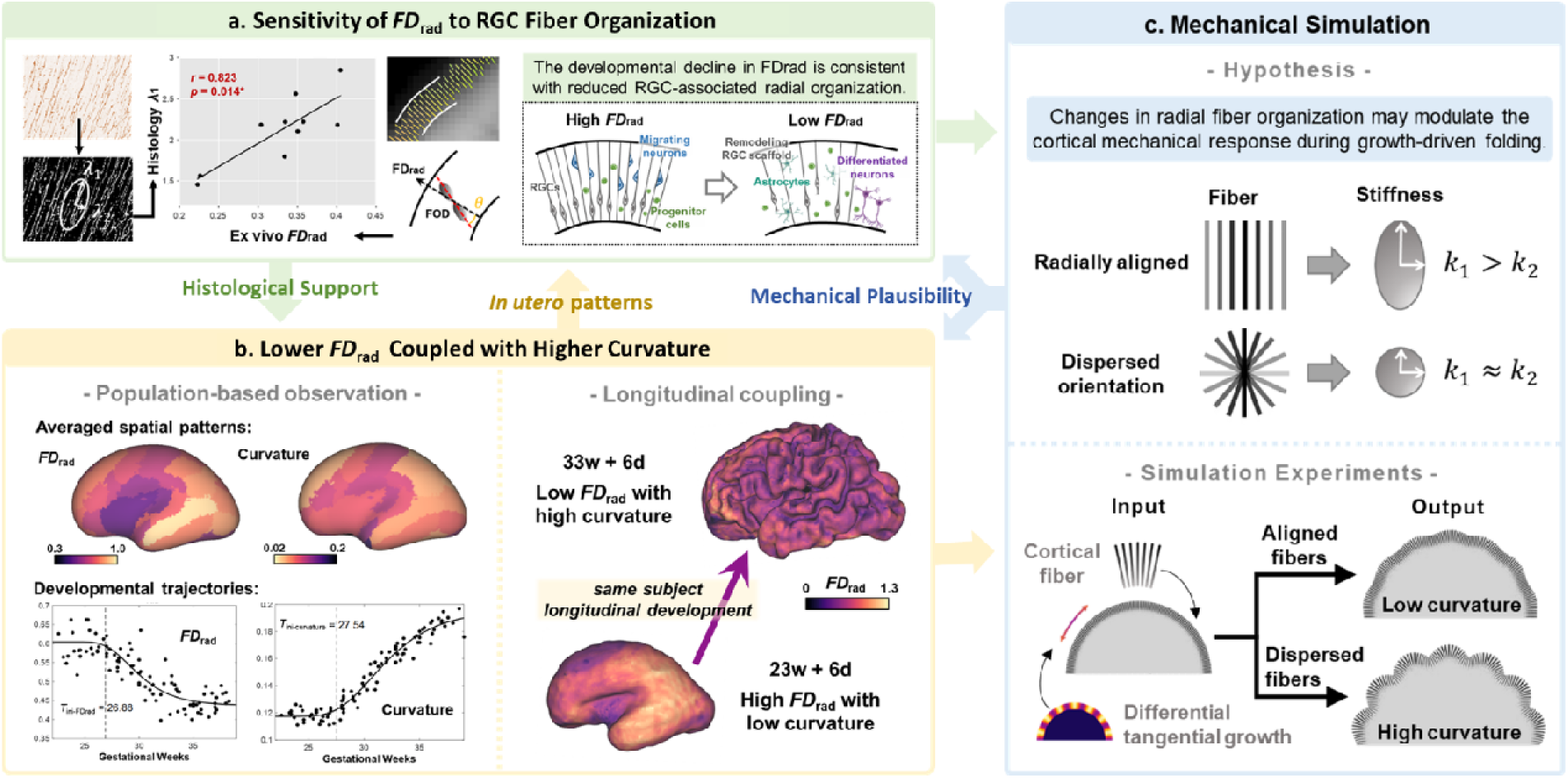
| Study Overview. **a**. Sensitivity of regional *FD*_rad_ to RGC-associated radial fiber organization. *Ex vivo* MRI and histological analyses showed the dMRI-based regional radial fiber density (*FD*_rad_) was positively associated with the principal structure-tensor magnitude derived from vimentin-labelled radial architecture, supporting its sensitivity to RGC-associated radial fiber organization. **b. Lower *FD*_rad_ is coupled with higher curvature.** Fetal cortical *FD*_rad_ and curvature showed linked spatial distributions. Cross-sectional and longitudinal in utero MRI consistently showed that reductions in cortical *FD*_rad_ were accompanied by increases in curvature, supporting coupled microstructural and morphological development (w, weeks; d, days). **c. Biomechanical simulation.** Computational mechanical simulations tested whether changes in modeled radial fiber content and orientation dispersion could modulate the cortical response to prescribed differential tangential growth. Under the model assumptions, reducing radial fiber support altered growth-driven buckling and produced greater cortical curvature under otherwise matched conditions, supporting the mechanical plausibility of this relationship.

**Fig. 2.**
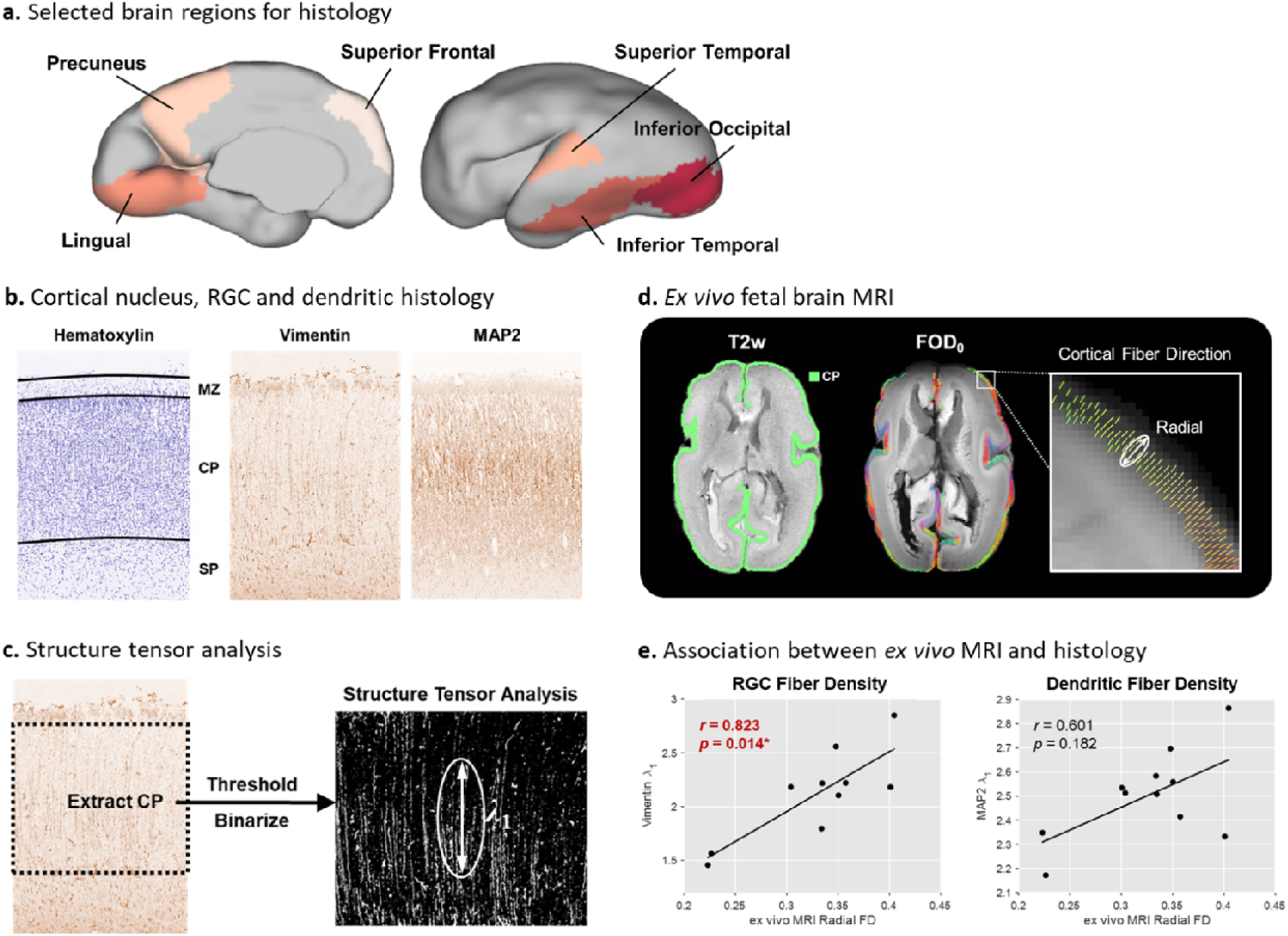
| *Ex vivo* MRI and immunohistology. **a**. Selected brain regions for histology. Representative brain regions in each lobe were extracted from the *ex vivo* brain sample. **b. Cortical nucleus, RGC, and dendritic histology,** as shown, labeled by Hematoxylin, Vimentin, and MAP2 staining. **c. Structure tensor analysis of RGC histology.** The dominant local structure-tensor response λ_1_ of Vimentin staining largely overlaps the radial fiber component. **d. *Ex vivo* MRI of the 23-week fetal brain sample.** The cortical plate (CP) was labeled on the T2w image, followed by cortical fixel-based analysis. FOD_0_: Map of the 0^th^-order spherical harmonic coefficient of the fiber orientation distribution (FOD). **e. Association between *ex vivo* MRI and histology.** Vimentin-derived λ_1_ was positively associated with *FD*_rad_ (*r* = 0.823, *P* = 0.014), while MAP2-derived λ_1_ was not significantly associated with *FD*_rad_.

MAP2 and vimentin both displayed positive staining in cortical regions (Fig. 2b), while Neurofilament staining was not evident in the examined cortical regions. Structural-tensor analysis was used to quantify the magnitude of the dominant locally oriented image structure within each immunostain (Fig. 2c). Quantitative analysis revealed that vimentin-derived λ_1_ was positively correlated with *ex vivo FD*_rad_ (*r* = 0.823, FDR-corrected *P* = 0.014) across 10 histologically examined brain regions (Fig. 2a). By contrast, the association between MAP2-derived λ_1_ and *ex vivo FD*_rad_ was not statistically significant (Fig. 2e). These findings demonstrate a regional correspondence between *FD*_rad_ and vimentin-labelled radial architecture within this specimen, supporting the sensitivity of *FD*_rad_ to RGC-associated radial fiber organization.

To determine whether the observed Vimentin association was also detectable using commonly used dMRI anisotropy measures, we additionally tested the correlations between vimentin-derived λ_1_ and other dMRI anisotropy indices, including FA, axial diffusivity (AD), and the maximum amplitude of fiber density (*FD*_max_). Notably, none of these alternative metrics exhibited significant correlations (FA: *r* = 0.562, FDR-corrected *P* = 0.118; AD: *r* = 0.526, FDR-corrected *P* = 0.118; *FD*_max_: *r* = 0.658, FDR-corrected *P* = 0.077). These findings underscore that *FD*_rad_, by explicitly incorporating the radial orientation prior, provides a specific and sensitive readout of the RGC architecture that standard dMRI markers fail to capture. Furthermore, none of the examined dMRI measures was significantly associated with the MAP2-derived structure-tensor measure (Supplementary Fig. S1).

### 2.2 *In utero* Developmental Patterns of Cortical *FD*_rad_ and Curvature

We then investigated the developmental trajectory of *FD*_rad_, as a measure of cortical radial fiber organization, in relation to cortical curvature. From 23 to 38 weeks of gestation, our *in utero* MRI dataset of the normal fetal brains (ZJU-Fetal, 131 pregnancies, including 91 T2w and 93 dMRI scans) ^12,33^ showed that the cortex undergoes significant morphological and microstructural changes simultaneously, as shown in Fig. 3. On T2w structural MRI, the majority of primary sulci were clearly visible starting from 27 weeks and grew deeper with accompanying secondary folds appearing. Prominent radially aligned fiber structures in the cortical plate around the primary folds were observed by dMRI before approximately 26 weeks and gradually diminished through gestation. Cortical fibers became more isotropically organized when approaching perinatal time.

The whole-cortex-averaged cortical curvature followed a Gompertz growth trajectory, with a model-derived timing parameter *T*_ini-curvature_ = 27.54 ± 0.31 weeks (*t* = 87.80, *P* < 0.001***). Conversely, the whole-cortex-averaged *FD*_rad_ followed an inverted Gompertz trajectory, with *T*_ini-*FD*rad_ = 26.88 ± 0.86 weeks (*t* = 31.15, *P* < 0.001***).

To further examine the directional component of the observed *FD*_rad_ trajectory, we performed a supplementary normalized-radiality analysis. For each cortical region, normalized radiality was calculated as regional *FD*_rad_ divided by the corresponding regional total apparent fiber-density measure, derived by summing apparent fiber density across fixels. Using the same modified four-parameter Gompertz model as for whole-cortex *FD*_rad_, normalized radiality showed a model-derived early transition location of 27.51 ± 0.25 weeks (95% CI, 27.02–28.00 weeks; *R*² = 0.961). It also remained negatively associated with curvature after adjustment for gestational age and regional effects (standardized β = −0.455, two-way cluster-robust *P* < 0.001). Together, these findings provide complementary evidence that the developmental decline in *FD*_rad_ reflects a reduction in radial organization relative to total apparent fiber density and is spatially coupled with cortical curvature (Supplementary Fig. S2).

### 2.3 Spatial Coupling between Cortical *FD*_rad_ and Curvature

Among the 78 cortical regions defined by the CRL atlas^34^, 52 regions met the original inclusion criteria for both measures: the Gompertz trajectories and their model-derived *T*_ini_ parameters were significant after FDR-correction (*P* < 0.05), and the estimated *T*_ini_ values fell within the observed gestational-age range. These regions were retained for the regional timing analysis. The *T*_ini-curvature_ and *T*_ini-*FD*rad_ maps showed similar spatial patterns, with the parietal and superior temporal lobes developing first (Fig. 4a). Pearson’s correlation revealed a significant positive correlation between *T*_ini-*FD*rad_ and *T*_ini-curvature_ across brain regions (*r* = 0.41, *P* < 0.01**). This correlation remained significant after accounting for cortical spatial autocorrelation using 10,000 surface-based rotations (two-sided *P*_spin_ = 0.0122), supporting the spatial coupling between the regional timing of cortical *FD*_rad_ reduction and curvature development (Fig. 4b).

**Fig. 3.**
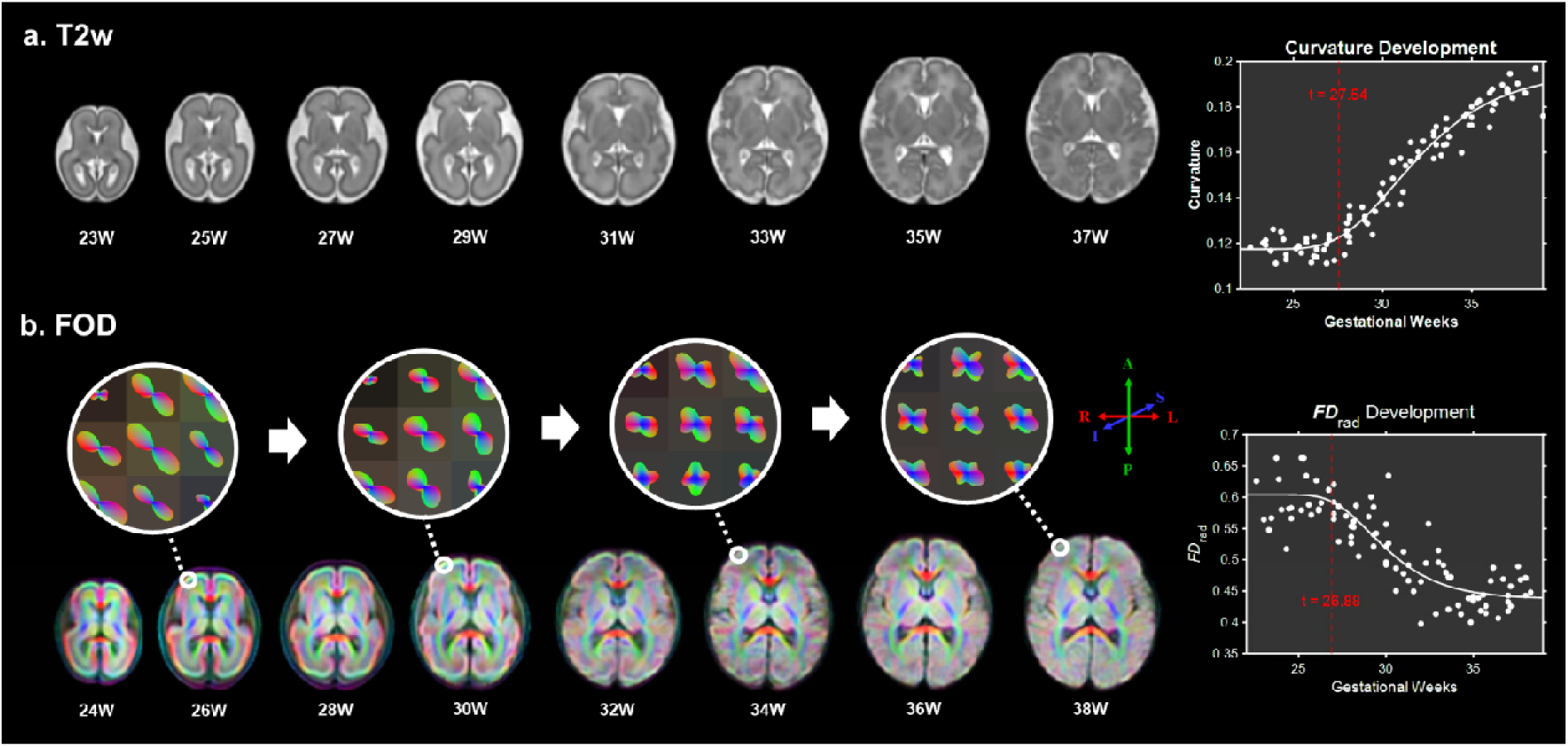
| *In utero* MRI of the fetal brain cortical radial fiber and curvature development. **a**. The T2w atlas of fetal brains was generated at each gestational week. The whole-brain-averaged cortical curvature followed a Gompertz growth curve, with a model-derived timing of curvature increase (*T*_ini-curvature_) at 27.54 ± 0.31 weeks (*t* = 87.80, *P* < 0.001***). b. FOD maps of the group-averaged fetal brains at each gestational week. Measured by the apparent fiber density along the cortical normal vector direction, the averaged whole cortical *FD*_rad_ followed an inverted Gompertz trajectory, with a model-derived starting time of decrease (*T*_ini-*FD*rad_) at 26.88 ± 0.86 weeks of gestation (*t* = 31.15, *P* < 0.001***).

**Fig. 4.**
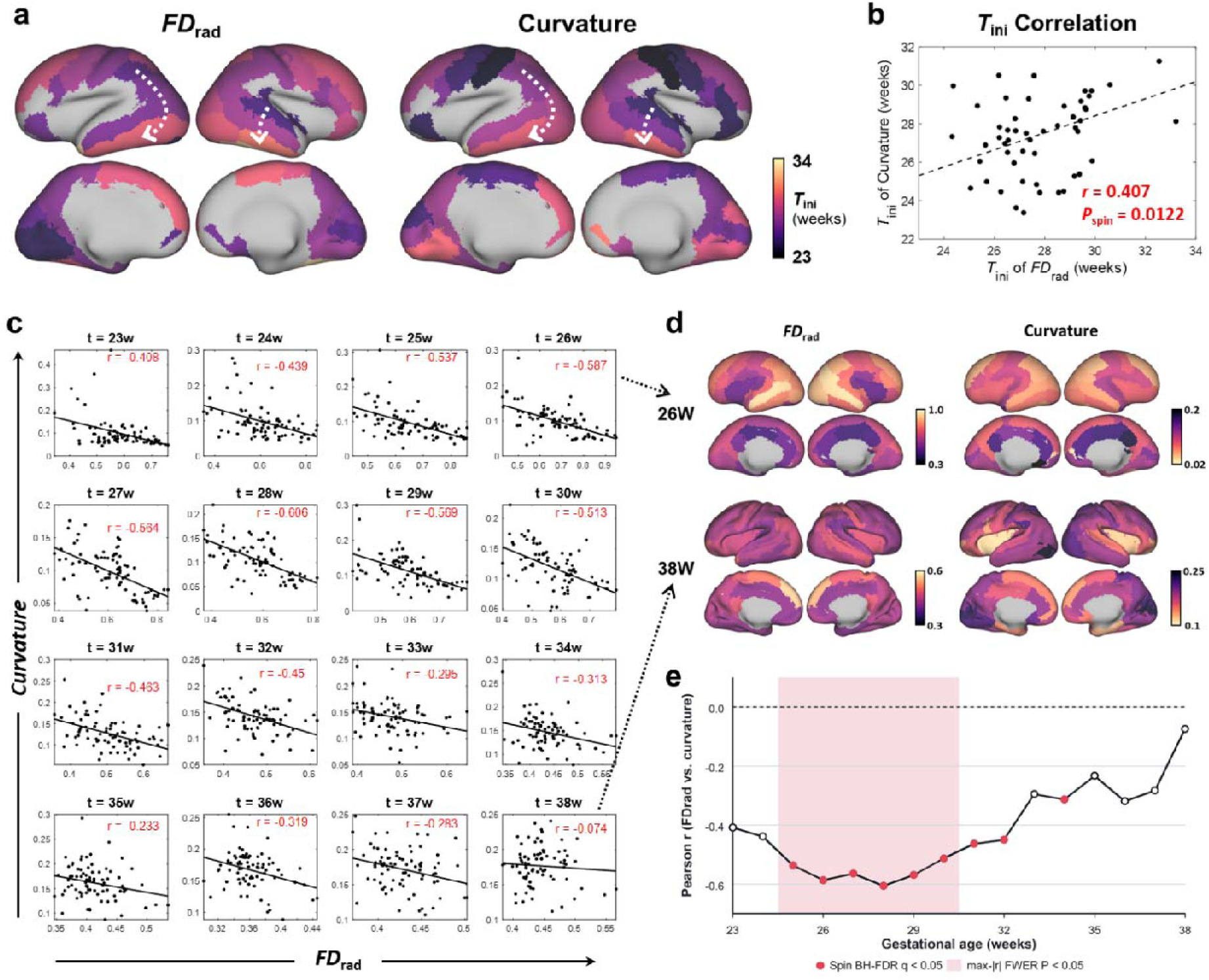
| Spatial association between *FD*_rad_ and curvature. **a**. The spatial pattern of *T*_ini_ on the CRL atlas-defined cortical region. The maps display the 52 regions in which both the FDrad and curvature trajectories and their model-derived *T*_ini_ parameters met the Gompertz-model inclusion criteria; unshown regions did not meet these criteria. *T*_ini_ had similar spatial patterns between *FD*_rad_ and curvature, with earlier model-derived timing of development in the parietal and superior temporal regions. **b. The correlation between *T*_ini-*FD*rad_ and *T*_ini-curvature_.**

Negative cross-regional correlations between *FD*_rad_ and curvature were observed from 23 to 38 weeks, reaching their greatest magnitude at 28 weeks and subsequently attenuating toward 38 weeks (Fig. 4c–e). After accounting for cortical spatial autocorrelation using surface-based spin tests and correcting across the 16 gestational weeks, the associations remained significant from 25 to 32 weeks and at 34 weeks (BH-corrected *q*_spin_ < 0.05). Under the more stringent max-|*r*| family-wise correction, the associations remained significant from 25 to 30 weeks (*P*_FWER_ < 0.05).

The regional *T*_ini_ maps were positively correlated after accounting for cortical spatial autocorrelation using 10,000 surface-based rotations (*r* = 0.407, two-sided *P*_spin_ = 0.0122). **c. Spatial correlation between *FD*_rad_ and curvature at each gestational week.** Negative cross-regional correlations were observed throughout the examined period, strengthening during mid-gestation, reaching their maximum at 28 weeks, and subsequently attenuating toward 38 weeks. **d. Exemplar spatial patterns of *FD*_rad_ and curvature.** The 26-week maps illustrate the pronounced negative spatial correspondence during mid-gestation, whereas this correspondence was weak at 38 weeks. The color bar for curvature was inverted for better comparison (w, weeks). **e. Developmental profile and spatial-null inference of the cross-regional *FD*_rad_–curvature correlation.** Red points indicate gestational weeks that remained significant after 10,000 surface-based rotations and BH correction across the 16 examined weeks (25–32 weeks and 34 weeks; *q*_spin_ < 0.05). The shaded interval indicates weeks that additionally survived the more stringent max-|r| family-wise error correction (25–30w; *P*_FWER_ < 0.05).

### 2.4 Coupled Longitudinal Changes in Cortical *FD*_rad_ and Curvature

The co-developmental pattern between cortical fiber organization and morphology was further examined using longitudinal *in utero* MRI. Utilizing an independent dataset of the developing Human Connectome Project (dHCP, https://biomedia.github.io/dHCP-release-notes/), we collected two longitudinal scans from the same subjects, one at mid-gestation (21.1−31.4 weeks) and one at late gestation (26.9−37.3 weeks). As shown in Fig. 5a, longitudinal changes in cortical *FD*_rad_ and curvature were consistent with the cross-sectional observation that decreasing *FD*_rad_ was accompanied by increasing curvature.

**Fig. 5.**
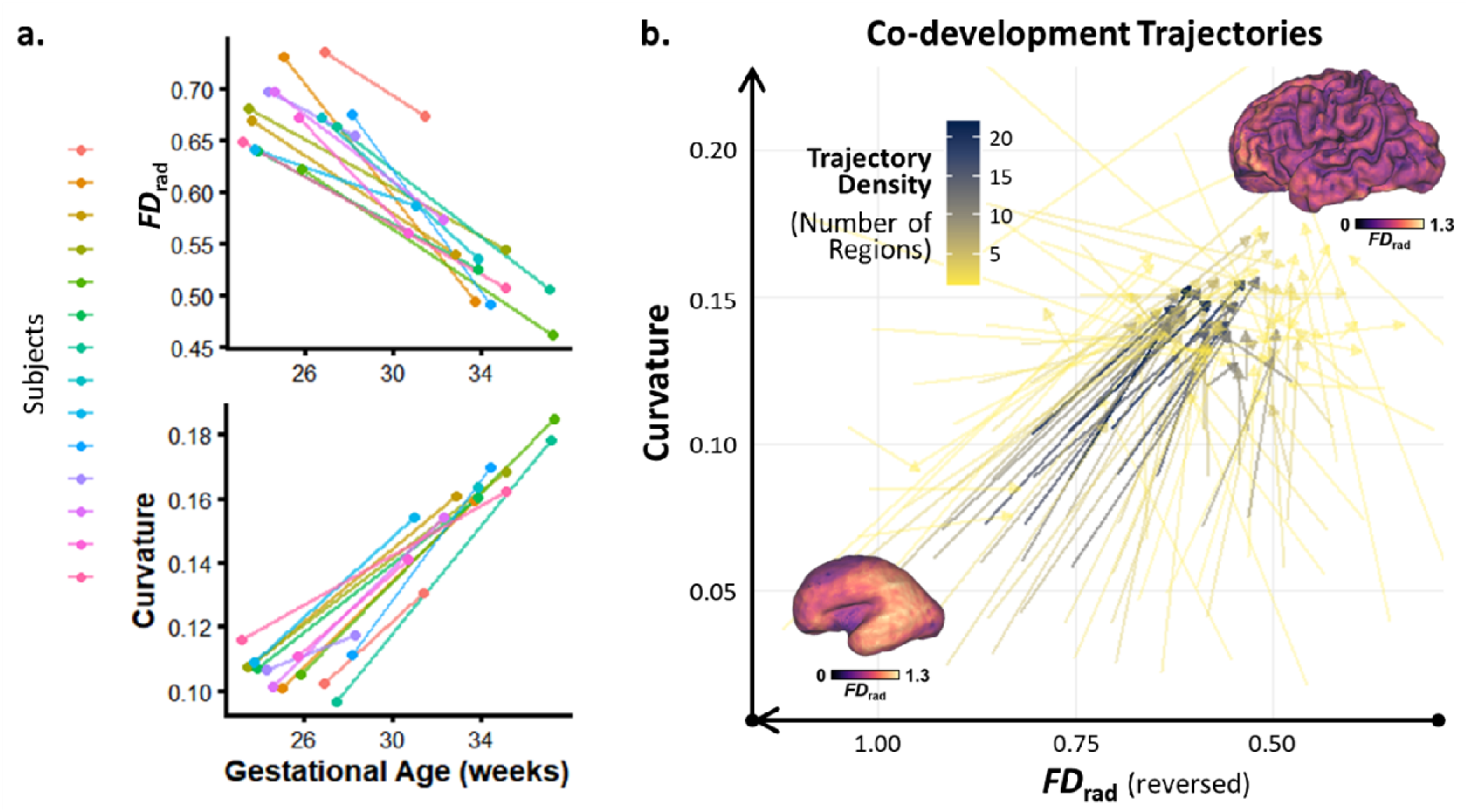
| Coupled longitudinal changes in cortical *FD*_rad_ and curvature. **a**. Longitudinal development from mid-to late-gestation. Longitudinal changes in cortical *FD*_rad_ and curvature were consistent with the cross-sectional observation that decreasing *FD*_rad_ was accompanied by increasing curvature. Only subjects within the studied GA range were displayed (GA > 23 weeks). **b. Co-developmental trajectories of regional *FD*_rad_ and curvature.** Individual trajectories from all subjects and cortical regions were aggregated based on path similarity. Each arrow represents a consensus co-developmental trend. The color intensity of the arrows indicates trajectory density (i.e., the number of regions across all subjects), with darker colors representing a higher frequency of occurrence. The “core development flow” summarized the predominant pattern from a high-*FD*_rad_/low-curvature state towards a low-*FD*_rad_/high-curvature state.

In the co-development linear mixed effect (LME) model, follow-up curvature was associated with baseline *FD*_rad_ (standardized β = –0.209, SE = 0.046, 95% CI [-0.298, –0.119], *z* = –4.55, *P* < 0.001) and the rate of *FD*_rad_ change (β = –0.280, SE = 0.041, 95% CI [-0.359, –0.200], *z* = –6.88, *P* < 0.001). Baseline curvature, included to adjust for the initial morphological state, was not significantly associated with follow-up curvature (β = –0.054, SE = 0.041, 95% CI [-0.135, 0.028], *z* = –1.30, *P* = 0.195). Follow-up curvature was also associated with baseline GA (β = 0.332, SE = 0.048, 95% CI [0.239, 0.426], *z* = 6.95, *P* < 0.001) and the interval between scans (β = 0.471, SE = 0.047, 95% CI [0.378, 0.563], *z* = 9.95, *P* < 0.001). The estimated variance components were 0.023 for subject, 0.278 for cortical region, and 0.353 for the residual. The phase-space plot of curvature and *FD*_rad_ change further illustrated their patterns of co-development (Fig. 5b). The “core development flow” originated from a high-*FD*_rad_/low-curvature state and progressed towards a low-*FD*_rad_/high-curvature state. Most longitudinal observations exhibited concurrent decreases in *FD*_rad_ and increases in curvature, visualized as long diagonal arrows pointing from the bottom-left toward the top-right of the phase plane. In addition, certain observations showed relatively little further reduction in *FD*_rad_ while curvature continued to increase, as indicated by the vertical upward-pointing arrows situated at the lower *FD*_rad_ levels (the right side of the plot). The principal co-developmental pattern was reproduced using the b = 400 s/mm² shell dataset (Supplementary Fig. S3).

### 2.5 Loss of Radial Fiber Alters Growth-Driven Folding in Mechanical Simulations

Next, we designed a computational model to examine the potential mechanical implications of our *in utero* observations. We tested whether reductions in radial fiber content or radial alignment altered the cortical mechanical response to differential tangential growth. The first two experiments examined: 1) how initial radial fiber density affected the extent of simulated folding; and 2) how the imposed timing of radial fiber loss affected curvature-growth timing.

In the first two experiments designed in this study, radial fiber density modulated the folding response under otherwise matched model conditions. First, lower initial radial fiber density in the cortex resulted in a greater extent of cortical folding. When the input fiber density was set to 1, 0.5, 0.1, and 0 per unit, the simulated curvature index at the endpoint of the simulation was 4.62, 5.06, 5.87, and 6.36 respectively (Fig. 6a). This quantitative trend was also reflected in the morphological evolution (Fig. 6b). Second, introducing the decrease in cortical radial fiber density earlier shifted curvature growth to an earlier simulation time. When the *T*_ini_ of fiber density (*T*_ini-fiber_) was set to 0s, 0.2s, 0.4s, and 0.8s of the simulation time, the resulting *T*_ini-curvature_ values were 0.83s, 0.87s, 0.90s, and 0.95s, respectively (Fig. 6a).

**Fig. 6.**
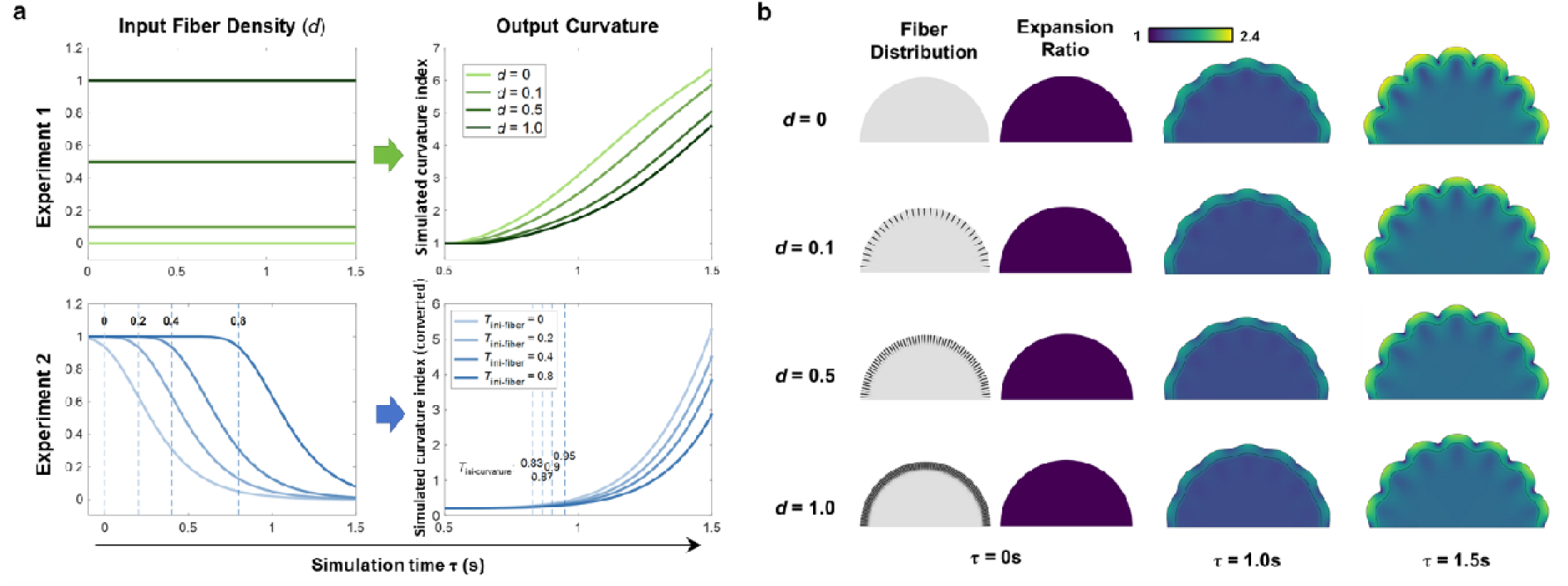
| Mechanical simulation results with different conditions of cortical fiber density. **a**. Computational experiments. Experiment 1: Lower initial radial fiber density in the cortex resulted in a higher curvature index. Experiment 2: An earlier prescribed decrease in cortical radial fiber density shifted the model-derived curvature growth to an earlier simulation time. **b. Model geometric evolution.** Before the formation of sulci, the stress induced by cortical surface expansion produced a modest increase in the diameter of the axisymmetric spherical model, shown in a meridional cross-section. After a period of continuous prescribed differential tangential growth, the cortical boundary progressively developed outward and inward deformations. Under otherwise matched growth conditions, models with higher initial radial fiber density exhibited lower curvature and less pronounced folding. The maps are colored according to the local tangential expansion ratio.

Furthermore, we adjusted fiber dispersion from strictly radial to a fully dispersed orientation distribution while holding total fiber density constant. This experiment represented progressive loss of coherent radial alignment within the same fiber population, rather than the addition of a separate isotropic fiber component, thereby distinguishing the contribution of fiber orientation from that of total fiber content. Specifically, we set the total fiber density to a fixed value of *d* = 1, and adjusted the fiber dispersion parameter κ from 0 (strictly radially aligned) to 0.33 (fully dispersed). As a result, simulated curvature increased more rapidly with increasing fiber dispersion. By a simulation time of 1.5 s, the simulated meridional curvature index increased from 3.99 in the radially aligned model to 5.87 in the fully dispersed model (Fig. 7a). Thus, even at a constant total fiber density, loss of coherent radial alignment altered the curvature response under identical growth conditions. Morphological evolution showed that the fully dispersed model developed deeper and more pronounced folds, whereas the radially aligned model exhibited a visibly smoother surface with restricted, shallow wrinkling (Fig. 7b).

**Fig. 7.**
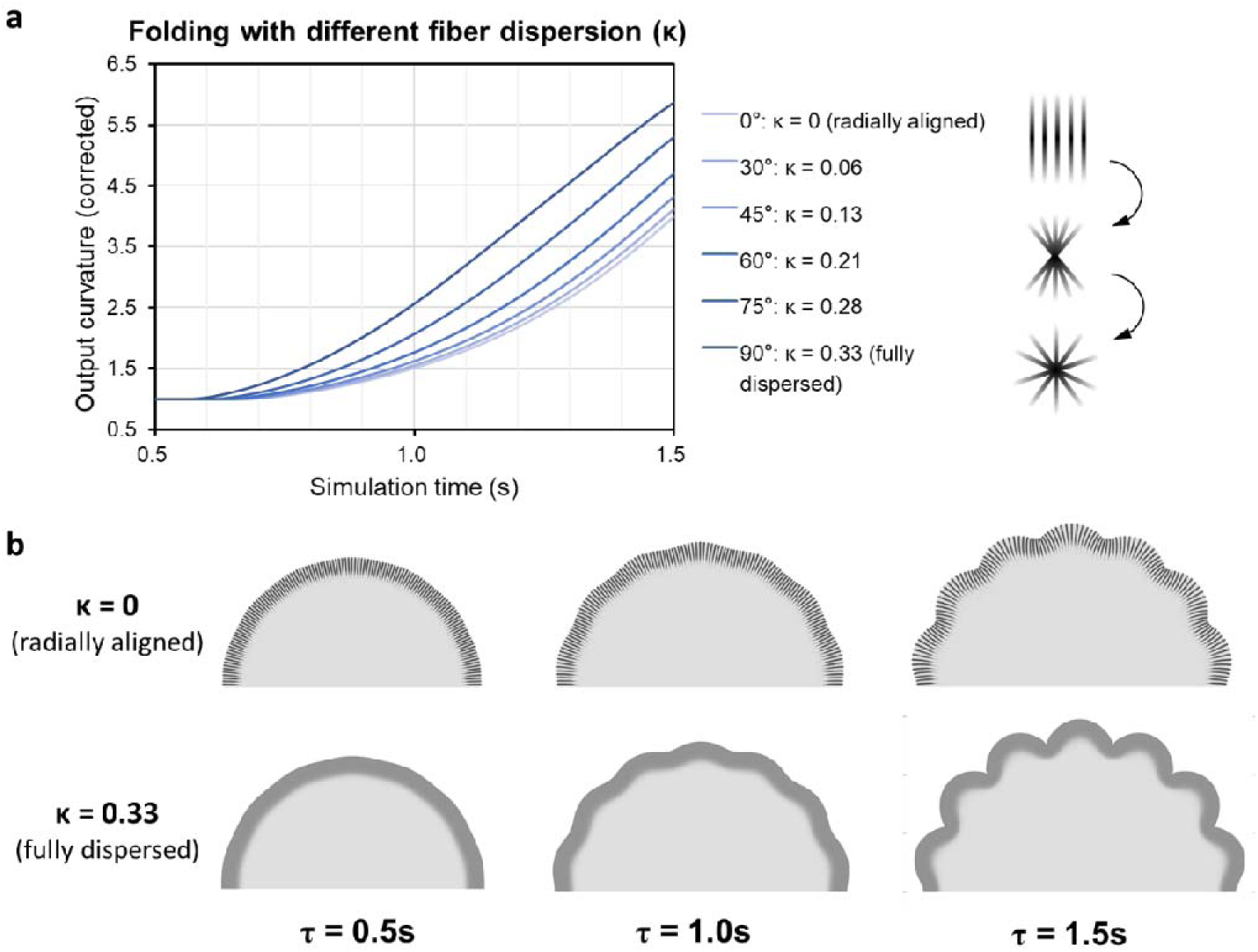
| Mechanical simulation results with different fiber dispersion. **a**. Relationship between fiber dispersion and simulated curvature development. Increasing the fiber dispersion parameter (κ) from strictly radial alignment (0°: κ = 0) toward a fully dispersed orientation distribution (90°: κ = 0.33) produced progressively faster curvature growth under otherwise matched model conditions. This result indicates that radial orientation coherence, independently of total fiber density, modulates the folding response. b. Morphological outcomes under different fiber-dispersion conditions. Simulation time points from 0.5s to 1.5s show the dynamic development. The fully dispersed model (κ = 0.33) exhibits deeper and more pronounced folds, while the radially aligned model (κ = 0) develops restricted, shallow wrinkling across a smoother topology.

## 3 Discussion

This study proposed diffusion MRI-based radial fiber density (*FD*_rad_) as a noninvasive measure sensitive to RGC-associated radial fiber organization. By combining *in utero* and *ex vivo* MRI, immunohistology, and computational modeling, our findings linked cortical microstructural reorganization with morphological development and examined its potential contribution to the biomechanical context of cortical folding. Comparison of *ex vivo* MRI with immunohistology showed that regional *FD*_rad_ was strongly associated with the principal structure-tensor magnitude derived from vimentin-labelled radial architecture, supporting its sensitivity to RGC-associated fiber organization. Our *in utero* MRI data further demonstrated spatially and temporally coordinated changes in cortical *FD*_rad_ and curvature, while longitudinal data from an independent dataset supported their coupled evolution within individuals. Finally, computational simulations showed that reductions in radial fiber content or alignment altered the cortical response to differential growth, supporting the mechanical plausibility of radial fiber remodeling as a modulator of growth-driven folding. Together, these complementary findings identify progressive loss of cortical radial organization as a prominent microstructural feature of the folding period and demonstrate its capacity to modulate growth-driven buckling within a mechanically plausible framework.

The relation between fiber organization and cortical folding was investigated previously. The classic “axonal pulling” theory hypothesized that the axonal tension pulls the connected brain regions inwards to form sulci^7^, although the extent to which this mechanism accounts for realistic fiber distributions and folding patterns remains debated^3^. Recent studies modeled subcortical fiber distribution as a result of brain folding-induced tension^4,35^. Other studies suggested that subcortical tension contributes to the patterns of folding rather than the morphogenetic process itself^5,6^. The current study introduces a complementary perspective by focusing on the cortical microstructure, especially radially organized fibers within the cortex, and examining how their assumed anisotropic mechanical contribution may modulate the pace of cortical folding. Our work therefore extends the differential-growth framework by identifying cortical cytoarchitecture as a developmentally varying mechanical modulator of folding. Within this framework, differential growth generates the compressive instability, while developmental loss of radial scaffold organization may modify the cortical mechanical response. This process likely overlaps with the development of subcortical white matter, axonal organization, neuronal differentiation, and other cellular events that collectively shape and stabilize the folding patterns^36^.

### 3.1 Cellular Substrate Underlying the Cortical Radial Architecture

The combined *ex vivo* MRI and histological analysis provided support for the biological basis of *FD*_rad_. Regional *FD*_rad_ was positively associated with the principal structure-tensor magnitude derived from vimentin-labelled radial architecture, supporting the sensitivity of *FD*_rad_ to RGC fiber organization. Although the dendritic component is undoubtedly a crucial element of cortical radial fibers, MAP2-derived structure-tensor measurements were not significantly associated with regional *FD*_rad_ in the examined 23-week specimen. A previous study reported that cortical FA value and folding patterns were associated with cortical dendrites^8^. That study examined a macaque brain in which primary folds had already formed and may therefore represent a later developmental stage. In the examined 23-week human fetal specimen, the stronger correspondence with vimentin-derived architecture identifies radial glial fibers as an important biological contributor to *FD*_rad_ during this developmental stage.

Together, the MRI and histological findings place the decline of the radial scaffold within the developmental period of rapid cortical folding. The tight relationship between the neuronal migration process along RGCs and cortical folding has been widely discussed, and the spatiotemporal patterns of neuronal migration are believed to set the stage for folding^37^. Aligning with our observations, Neal, et al.^38^ characterized the patterns of neurodevelopmental trajectories in relation to the onset of cortical convolutions in the ferret brain using MRI, demonstrating that gyrification becomes prominent after the completion of neuronal migration. Crucially, the regulation of cytoskeletal proteins is known to dictate the cellular structure and polarity of RGCs^30,39^. As the guiding function of the RGC scaffold declines, its reorganization may alter the radial architecture and anisotropic mechanical contribution of the cortical tissue. We propose that this change provides a more permissive mechanical context for tangential-expansion-driven buckling.

### 3.2 Spatiotemporal Coupling of Microstructural and Morphological Development

*In utero* MRI revealed coordinated development between cortical microstructure and morphology. The correspondence between the regional *T*_ini-*FD*rad_ and *T*_ini-curvature_ maps supports coordinated regional timing of radial-fiber decrease and curvature development. The coordination of microstructural and morphological development has been hinted at by previous studies; for instance, Takahashi, et al.^40^ observed that multi-oriented cortical fibers emerged before the appearance of primary sulci. Other studies have reported a decrease in fractional anisotropy^13,41^, and loss of radial coherence in the developing fetal brain cortex^42,43^, aligning with our quantitative observation of radiality loss.

Notably, the negative spatial association between *FD*_rad_ and curvature was most pronounced during mid-gestation and attenuated toward late gestation. This developmental profile suggests that the relationship between radial organization and cortical morphology changes across stages of gyrification. The phase-space trajectories (Fig. 5b) similarly showed continued curvature increase after *FD*_rad_ changes became smaller, consistent with stage-dependent coupling.

The independent longitudinal analysis extended the cross-sectional findings to within-individual development, showing that reductions in *FD*_rad_ and increases in curvature were coupled over the same developmental interval. Regarding the upstream molecular programs that drive this microstructural priming, our previous study linking *in utero* MRI with gene expression revealed that SYNE2, a cortical curvature-related gene, is significantly enriched in the biological process of neuronal migration along RGCs^21^. This provides convergent molecular context for the developmental link between RGC scaffold dynamics and cortical folding. Taken together, the regional, gestational-age-resolved, and within-subject findings provide convergent evidence that progressive loss of cortical radial organization is closely coupled to the emergence and elaboration of cortical folds, identifying the radial scaffold as a biologically grounded candidate modulator of cortical morphogenesis.

### 3.3 Reduced Radial Fiber as a Potential Modulator of Cortical Folding

Our biomechanical simulations demonstrated that changes in radial fiber organization can modulate the pace and extent of growth-driven folding within the model. Differential growth generated the compressive instability, whereas radial fibers supplied anisotropic reinforcement that resisted buckling. Reducing fiber content or radial coherence consistently increased curvature under matched growth conditions, demonstrating that radial organization exerted a mechanically meaningful influence on the folding response.

Therefore, our simulation was designed to go beyond static anatomical replication to capture how the evolving microstructure influences the dynamic trajectory of morphogenesis. The fiber-density experiments showed that changing the timing of radial-fiber reduction shifted the timing and pace of the simulated folding response. The dispersion experiment further showed that radial orientation coherence influenced folding even when total modeled fiber density was held constant, indicating that fiber organization as well as content can shape the simulated mechanical response.

The relation between mechanical properties, particularly stiffness, and morphological folding has been explored in several studies. Specifically, simulation studies have demonstrated that stiffness distribution dictates the wavelength of cortical folding patterns^44,45^. Furthermore, Zarzor, et al.^46^ highlighted the importance of incorporating dynamic changes in local tissue stiffness derived from cell density, while others have argued that bulk tissue stiffness results from the combined effects of cellular architecture and existing mechanical forces rather than individual cellular properties alone^25^. By holding growth and other material parameters constant, our simulations isolated the contribution of radial anisotropy to the folding response. Thus, differential growth supplies the compressive drive, while progressive loss of the radial scaffold increases the cortex’s mechanical susceptibility to buckling, providing a complementary link between cellular reorganization and tissue-scale morphogenesis.

### 3.4 Limitations

Several limitations of this study should be acknowledged. First, the observational MRI analyses establish spatial and longitudinal coupling between cortical radial organization and curvature but do not independently determine causal direction or a universal temporal sequence. The simulations complement these observations by testing a mechanically plausible modulatory mechanism within a controlled framework, rather than identifying a unique biological cause of cortical folding.

Second, the histological analysis was based on one 23-week human fetal brain. Given the exceptional rarity of well-preserved fetal specimens with matched ex vivo MRI and histology, the region-wise analysis across 10 anatomically distributed cortical areas provided valuable within-brain histological support for the sensitivity of FDrad to vimentin-labelled radial architecture. Replication in additional specimens and developmental stages will be important for establishing population-level generalizability and directly characterizing the histological trajectory of radial-scaffold disassembly.

Third, the simulations were designed as controlled mechanistic experiments rather than subject-specific reconstructions of fetal cortical development. Holding growth and other material parameters constant allowed the contributions of radial fiber content and orientation coherence to be examined independently. However, *FD*_rad_ was not quantitatively mapped to tissue stiffness, and simulation time was not mapped to gestational age. Future anatomically realistic models should incorporate heterogeneous cortical and subcortical properties, the continuous radial scaffold across developmental compartments, a broader range of fiber organizations, and quantitative MRI-to-model mapping.

## 4 Methods

### 4.1 The Fetal Brain MRI Datasets

#### 4.1.1 Cross-sectional *in utero* MRI

The ZJU-Fetal dataset included 131 pregnant women aged between 18 and 42 years (mean age = 30.1 ± 4.8 years). The modality-specific analyses included 91 fetal brain T2-weighted scans from 89 participants and 93 fetal brain diffusion MRI scans from 92 participants. Fifty-one women contributed both modalities, yielding 52 paired T2w–dMRI sessions. Two participants underwent repeat T2w imaging, and one of them also underwent repeat dMRI, contributing two paired sessions. Among the 130 participants with at least one analyzable modality, 41 contributed dMRI only and 38 contributed T2w MRI only. The detailed acquisition protocol can be found in papers by Xu, et al.^33^ and Chen, et al.^12^. In addition, our previously published spatiotemporal atlases of normative fetal brain development, including both T2w (https://github.com/zjuwulab/CHN-fetal-brain-atlas) and dMRI (https://github.com/zjuwulab/Fetal-Brain-dMRI-Atlas) data, were incorporated to provide standardized reference information^12,33^. Quantitative cortical measurements were extracted from both individual scans and atlas-space representations to delineate developmental trajectories and to inform mechanical simulations. The in utero fetal MRI study was approved by the Ethics Committee of Shandong Provincial Hospital (NSFC:NO.2019-062). Written informed consent was obtained from all participants.

#### 4.1.2 Longitudinal *in utero* MRI

Longitudinal fetal brain MRIs were obtained from the Developing Human Connectome Project (dHCP) fourth data release, and 16 subjects were selected with two longitudinal scans. GA at first scan ranged from 21 weeks + 1 day to 31 weeks + 3 days (mean GA = 25.06 ± 2.38 weeks), and GA at the second scan ranged from 26 weeks + 6 days to 37 weeks + 2 days (mean GA = 32.57 ± 3.02 weeks). At each time point, T2w structural MRI and dMRI were acquired. The *in utero* imaging protocol and the data processing details of the longitudinal dataset can be found in the notes for the dHCP data release (https://biomedia.github.io/dHCP-release-notes/).

#### 4.1.3 *Ex vivo* MRI

To examine the histological basis of the MRI–derived radial organization measure, an *ex vivo* fetal brain sample at 23 weeks was collected from the Women’s Hospital of Zhejiang University (WHZJU). The procurement and research use of the *ex vivo* fetal brain specimen were approved by the Ethics Committee of WHZJU (approval no. IRB-20230274-R). Written informed consent for tissue donation and research use was obtained from the parents before specimen collection. The whole brain was fixed in 10% formalin for over 3 months before MRI. The sample was then washed using normal saline solution for 1 week and in Fomblin oil during the MRI scan. *Ex vivo* MRI was performed on a 7.0 T Bruker BioSpin MRI scanner. The T2w image was acquired using the T2 Turbo RARE sequence, with TR = 12,878 ms, effective TE = 62.2 ms, in-plane resolution = 0.25×0.25 mm², and slice thickness = 0.5 mm. dMRI was acquired using a 2D SE EPI sequence, with TR = 6000 ms, effective TE = 28.07 ms, in-plane resolution = 0.25×0.25 mm², slice thickness = 0.6 mm, slice gap = 0.2 mm, three b = 0 images and 30 diffusion-weighted directions at b = 4000 s/mm^2^. Note that due to the substantially reduced water diffusivity in *ex vivo* samples compared to *in vivo*, higher b-values are required for *ex vivo* dMRI^47^.

### 4.2 MRI Data Processing

Image preprocessing strategies were adapted to the specific acquisition protocols of each dataset. For the in-house i*n utero* dataset, T2w MRI images underwent slice-to-volume registration (SVR) and super-resolution reconstruction (SR) using NiftyMIC. Corresponding dMRI images were SVR-SR reconstructed using SVRTK^48^. The preprocessed subject images were spatially normalized to a population-based weekly fetal brain atlas. Detailed image processing pipelines were described in our previous works^12,33^. For the dHCP fetal dataset, pre-processed data were utilized as per the official release notes. Specifically, T2w images were processed via the dHCP structural pipeline, while dMRI data underwent the SHARD reconstruction pipeline^49^.

Following reconstruction, a unified downstream analysis framework was applied to all three datasets. Cortical surfaces and vertex-wise mean curvature were derived using the dHCP structural pipeline. For the *ex vivo* T2w image, the same pipeline was applied after manual refinement of the cortical tissue labels. At each vertex of the white-matter surface, mean curvature was defined as the arithmetic mean of the maximum and minimum principal curvatures, which characterize surface bending along two mutually orthogonal tangential directions. The absolute value was taken at each vertex before regional averaging, thereby preventing cancellation between convex and concave curvature. Vertex-wise curvature was mapped to the image grid using Connectome Workbench metric-to-volume mapping with nearest-vertex assignment.

For the dHCP dataset, single-shell FOD estimation was adopted to match the acquisition framework of the in-house cross-sectional data. The b = 1000 s/mm² shell was selected for the primary longitudinal analysis because its stronger diffusion weighting provides greater sensitivity to directionally restricted tissue components while retaining adequate signal-to-noise ratio in the reconstructed fetal dMRI data. The b = 400 s/mm² shell was analyzed independently as a lower-diffusion-weighting qualitative robustness analysis.

Fiber orientation distributions (FODs) and fixel-based analysis (FBA) were subsequently generated using MRtrix3 to obtain fixel-wise apparent fiber density and corresponding fixel orientations^50^. The FOD data and derived fixel orientations were brought into T2w image space using the same linear registration. Vertex-wise normal vectors were calculated from the reconstructed mid-thickness surface and mapped to the registered dMRI voxel grid using nearest-vertex metric-to-volume mapping. The mapped surface normals and fixel orientations were therefore represented in the same image coordinate system. Each cortical voxel contained one mapped surface-normal vector and one or more fixel orientations, enabling voxel-wise calculation of the alignment between every fixel axis and the local cortical normal. Surface-normal vectors were renormalized to unit length before angle calculation.

To specifically quantify the radially organized cortical fibers, we designed a diffusion-derived measure termed radial fiber density (*FD*_rad_), calculated by weighting fixel-wise apparent fiber density according to its alignment with the local cortical surface normal:

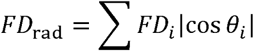

where *FD_i_*was the fiber density along the *i*th fixel within each voxel, and was the angle between the *i*th fixel direction *v*_fixel,*i*_ and the surface normal vector *v*_surface_, formulated as

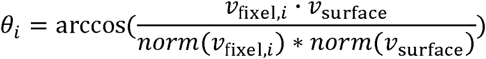

The absolute cosine was used because fixels represent unoriented axes. Thus, opposite vectors along the same anatomical axis contributed equally to *FD*_rad_.

As a supplementary sensitivity analysis, normalized radiality was calculated for each scan and cortical ROI as regional *FD*_rad_ divided by the corresponding regional total apparent fiber-density measure, derived by summing apparent fiber density across all fixels using the same cortical mask and regional mapping. Full procedures are provided in the Supplementary Materials.

The CRL fetal brain atlas was utilized to define the 78 cortical parcellations. Individual surfaces were aligned to the weekly surface templates using Multimodal Surface Matching (MSM)^51^. Mean absolute curvature and *FD*_rad_ values were extracted for all *in utero* cortical regions, while *ex vivo FD*_rad_ was extracted for the histologically examined regions.

### 4.3 Fetal Brain Histology

#### 4.3.1 Tissue Preparation

Fetal brain histology was performed after MRI. The whole brain was retrieved and immersed in normal saline solution to wash off the remaining Fomblin oil. Representative brain regions from each lobe were sampled by an experienced histologist, including the bilateral medial superior frontal gyrus, orbital frontal cortex, superior and inferior temporal gyri, precuneus, lingual gyrus, and inferior occipital lobe. Each tissue block was paraffin-embedded and sectioned at 4 µm thickness in a plane perpendicular to the cortical surface. The paraffin-embedded tissue sections were then deparaffinized and rehydrated for immunohistology.

#### 4.3.2 Pilot Experiment

Before the formal staining experiment, HE staining was initially performed to assess the quality of the preserved tissue and select target regions. For the selected regions, a pilot test was performed for each immunohistochemistry item to determine the optimal antibody dilutions. For anti-MAP2, anti-NF-L, and anti-vimentin antibodies, three dilutions of 1:2000, 1:4000, and 1:8000 were tested. Appropriate positive controls were included to ensure technical validity. An optimal dilution was selected based on visual examination by two experienced histologists. The staining procedures are described below.

#### 4.3.3 Immunohistochemistry

To avoid technical discrepancies, all regional tissue blocks for each immunohistochemistry target were processed in the same batch. Briefly, sections were immunolabeled using primary antibodies against Neurofilament-L (axons; Abcam, ab223343), MAP2 (dendrites; Abcam, ab254264), and Vimentin (RGCs; Abways, CY5134). Following overnight incubation at 4 °C, sections were treated with an HRP-conjugated secondary antibody and visualized using DAB.

Following the DAB chromogenic reaction, sections were rinsed with tap water and counterstained with hematoxylin, differentiated in acid alcohol, and blued in an alkaline solution. Finally, the slides were dehydrated through graded ethanol, cleared in xylene, and mounted with a permanent mounting medium. The stained slides were examined under a bright-field microscope and digitized using a whole-slide scanner for further analysis.

#### 4.3.4 Histological Data Processing

Multi-channel histological images were preprocessed using QuPath-0.6.0 to perform color deconvolution to separate the DAB channel (the target stain) and the Hematoxylin channel (the nuclei)^52^. A cortical ROI was segmented on the Hematoxylin channel image using intensity-based thresholding with subsequent manual refinement, extracting a contiguous cortex section. The DAB channel underlying the corresponding area was then sent to ImageJ for further analysis^53^. The greyscale image was first auto-thresholded by ImageJ and binarized to remove background signal, preserving the positively stained cells only.

Structure tensor analysis of the resulting binary images was performed using MATLAB R2021a:

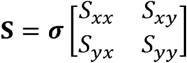

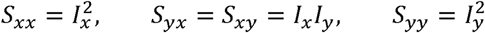

where a Gaussian kernel σ was adopted to downsample histology to a comparable resolution to *ex vivo* MRI. *I_x_* and *I_y_* were the image gradients. The eigenvalues of the structure tensor were:

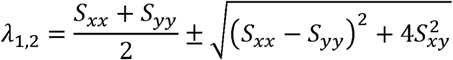

The principal eigenvalue λ_1_ quantifies the magnitude of the dominant local structure-tensor response. In the 23-week specimen, the dominant stained structures were predominantly radial. Mean λ_1_ was calculated within each histological ROI as a summary measure of the strength of locally organized stained architecture.

To assess the regional correspondence between histological architecture and diffusion-derived measures, Pearson’s correlation analysis was performed between vimentin– and MAP2-derived λ_1_ and *ex vivo* MRI-based *FD*_rad_, *FD*_max_, FA, and AD. An FDR-corrected *P* < 0.05 was considered statistically significant.

### 4.4 Computational Mechanical Simulation

#### 4.4.1 Model of cortical folding

We designed an axisymmetric finite-element model defined on a two-dimensional meridional semicircular cross-section of an initially spherical geometry. Under the axisymmetric assumption, this cross-section represents the corresponding three-dimensional body of revolution, while the three-dimensional constitutive and growth formulations, including the circumferential component, were retained. Simulation results are presented as two-dimensional meridional cross-sections for clarity.

The elastic response of brain tissue was modeled using a neo-Hookean material^3,4^, whose strain energy density function was:

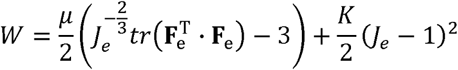

where **F_e_**was the elastic deformation tensor and *J*_e_ = det(**F**_e_), accounting for the elastic deformation-induced regional volumetric change. In previous studies, the shear modulus μ of fetal brain tissue has generally been set to values ranging from tens to hundreds of Pascals (Pa), with bulk modulus *K* significantly larger than μ^3,4^. However, the precise physiological value remains elusive due to the inherent difficulties of *in utero* measurement. In this context, these values were defined as μ = 100 Pa and *K* = 100μ.

In continuum mechanics, the total deformation can be formulated as **F = F_e_** · **G**, where **G** is the growth tensor **G =** [*G*_1_, *G*_2_, *G*_3_]*^T^*, with both tangential and radial growth components. In our model, the cortical plate (CP) and subcortical tissue were modeled as separate compartments within a unit sphere (*r* = 1) (Figure 8). Specifically, the CP was defined as the outer region occupying the radial interval from *r* = 0.9 to *r* = 1.0. In the CP, radial growth was set to unity because the fetal brain thickness changes little, while tangential growth was defined as an exponential function based on model fitting for *in utero* measurements in our previous study^33^:

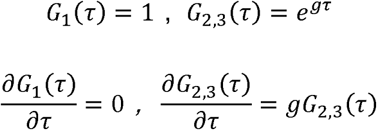

where τ was the simulation time. The tangential growth rate *g* was set to 0.01. A small spatial perturbation, *g\** = 0.1*g*(cos7θ)^2^, was superimposed on the tangential growth field to break spatial homogeneity and seed reproducible fold locations. The same perturbation was applied in all simulation experiments and was independent of fiber density and orientation dispersion. No independent prescribed growth was assigned to the subcortical compartment. Its evolution was governed by the specified stress-dependent growth formulation:

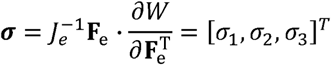

**Fig. 8.**
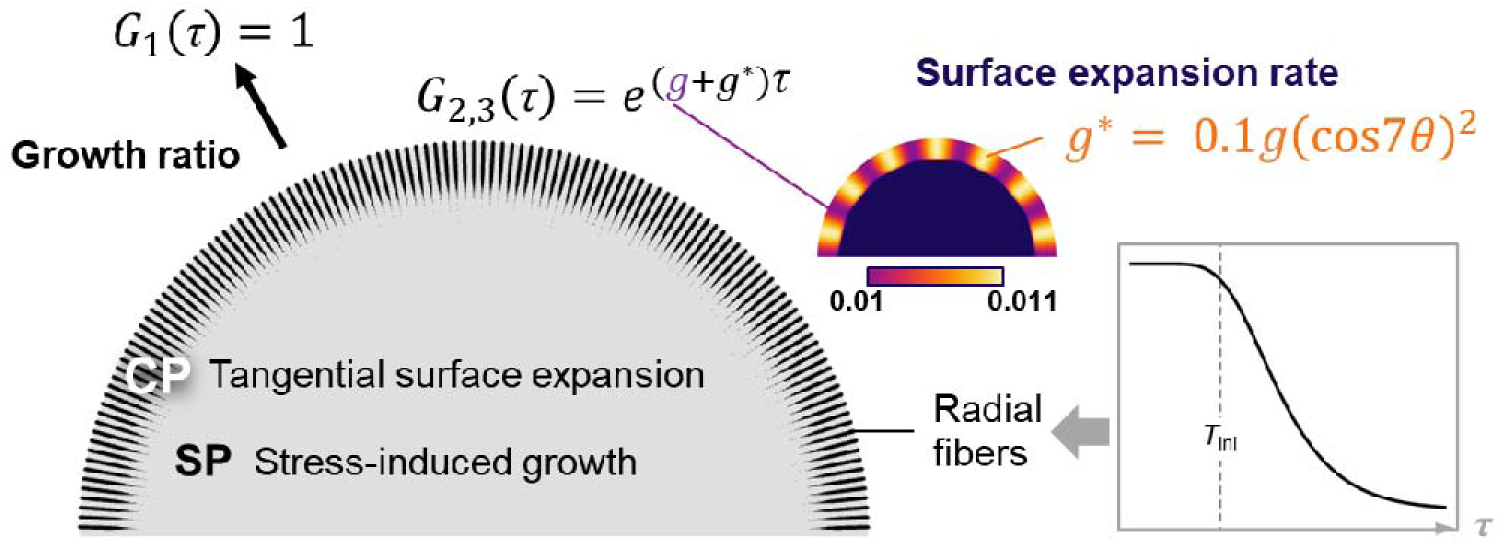
| Computational simulation model design. We designed an axisymmetric spherical model represented by a two-dimensional meridional semicircular cross-section as an initial morphology. The CP and subcortical tissue were defined as separate compartments. A regional difference *g\** = 0.1*g*(cos7θ)^2^ was introduced in the CP expansion to allow symmetry breaking. Radial fibers were embedded as anisotropic Holzapfel-Gasser-Ogden materials within the CP compartment. Fiber density and orientation dispersion were varied in the simulation experiments.

The subcortical tissue growth ratio was

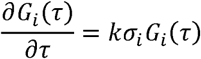

where *k* is the stress-induced growth rate. The stress-induced growth coefficient was set to k = 0.006 following preliminary numerical calibration to obtain stable subcortical deformation and a morphologically interpretable folding trajectory over the simulated interval. This value was held constant across all experiments, ensuring that between-condition differences arose from the manipulated radial-fiber parameters rather than changes in subcortical growth. The final problem to be solved can be formulated as:

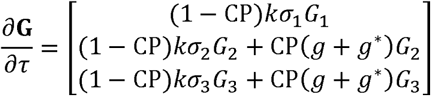

This formula describes the instantaneous growth ratio related to both its volumetric growth and stress-induced deformation.

#### 4.4.2 Cortical fiber modeling

Previous mechanical studies have shown that anisotropically organized neuronal fibers exhibit anisotropic stiffness. More specifically, higher stiffness was observed along the axial direction of fibers, while no difference in the direction perpendicular to the fibers was observed compared to the surrounding tissue^54,55^. Because the mechanical properties of fetal radial glial fibers have not been measured directly, their contribution was represented as a constitutive assumption rather than a parameter estimated from MRI. Accordingly, cortical tissue containing modeled radial fibers was represented using the anisotropic Holzapfel-Gasser-Ogden (HGO) materials within the CP compartment^55^. The anisotropic fiber contribution was activated predominantly under tensile stretch, consistent with the tension-bearing behavior observed in aligned neural processes and with the standard tension–compression assumption of distributed-fiber constitutive models ^3,54,55^. Because the compressive response of fetal RGC processes has not been measured directly, this treatment was adopted as a constitutive idealization. The fiber-associated stiffness parameter was set to 3μ for the controlled simulation experiments. This value was not quantitatively calibrated to *FD*_rad_ or measured fetal tissue stiffness.

Although RGC processes extend continuously from the ventricular zone through the intermediate zone and subplate to the pial surface, the anisotropic fiber term was restricted to the CP because both *FD*_rad_ and curvature were quantified in this compartment and the simulations were designed to isolate the local cortical contribution of radial organization. The deeper portions of the scaffold were subsumed into the effective constitutive response of the subcortical compartment rather than modeled as a separate translaminar fiber network.

We designed three computational experiments to investigate how radial fiber density, the timing of fiber-density reduction, and fiber orientation dispersion modulated differential-growth-driven cortical folding. For each simulation, time-dependent equations were solved, and folding was quantified along the boundary between the CP and subcortical compartments, corresponding to the white-matter surface used for MRI-based curvature estimation. A weak-form boundary PDE was used to obtain continuous boundary-normal components, and the local meridional curvature index was calculated as *C*_sim_ = |*w*_1_*T*_r_ + *w*_2_*T*_z_|/2, where *w*_1_ and *w*_2_ represent the radial and axial components of the interface normal vector, respectively, and *w*_1_*T*_r_ and *w*_2_*T*_z_ describe their changes along the interface. Their sum quantifies bending within the meridional cross-section. The absolute value removes dependence on normal-vector orientation, and the factor of 1/2 follows the definition of the curvature index used consistently across simulations. The line-averaged *C*_sim_ was recorded at each simulation time point and used to compare folding trajectories across experimental conditions. This simulated meridional curvature index is conceptually related to, but not numerically equivalent to, the three-dimensional mean absolute surface curvature derived from MRI.

##### Experiment 1: Static Fiber Density

The first experiment aimed to determine whether the magnitude of modeled radial fiber density altered the pace and extent of growth-driven cortical folding under otherwise matched conditions. To this end, the embedded relative fiber density (*d*) was controlled at static values. Four independent simulations were carried out with *d* set to 1, 0.5, 0.1, and 0, respectively.

##### Experiment 2: Prescribed Dynamic Reduction in Fiber Density

The second experiment investigated whether introducing fiber-density reduction at different simulation times shifted the model-derived curvature-change time. Fiber density was modeled as a time-dependent variable that declined from *d* = 1 to *d* = 0. This manipulation represented a generic loss of modeled radial fibers. It was governed by a reversed Gompertz growth function:

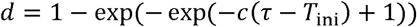

where *c* = 5 was defined to match the total simulation time. *T*_ini_ was set to 0 s, 0.2 s, 0.4 s, and 0.8 s. Note that simulation time τ represents the time steps of the evolving model and does not directly correspond to gestational age. The output curvature was fitted to a Gompertz growth model to estimate its model-derived time parameter.

##### Experiment 3: Fiber Dispersion and Radial Orientation Coherence

While the first two experiments maintained strictly radially aligned fibers (κ = 0), the third experiment was designed to explicitly isolate the specific contribution of fiber dispersion to cortical folding. According to the HGO material property, the mechanical response of a dispersed fiber orientation distribution to deformation is characterized by a modified pseudo-invariant *I*_ak_, formulated as

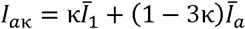

Where 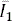 captures the isotropic bulk distortion, and 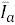 represents the anisotropic stretch along the fiber direction. At κ = 1/3 ≈ 0.33, the modeled orientation distribution reaches the fully dispersed limit. This condition does not represent the addition of a separate population of isotropic fibers. The correspondence between the parameter κ and the physical dispersion half-cone angle θ is 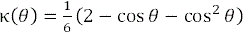. In this experiment, total fiber density was held constant (*d* = 1), and we compared radially aligned models against dispersed models with θ ranging from 30° to 90° in 15° increments, corresponding to κ □ {0.06, 0.13, 0.21, 0.28, 0.33}.

### 4.5 Statistical Analysis

We used a modified Gompertz growth model to fit the developmental trend of *FD*_rad_ and cortical curvature with gestational age, which is formulated as:

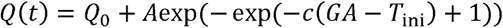

where *Q*_0_ denotes the fitted baseline, *A* the signed trajectory amplitude, *c* the rate parameter, and *T*_ini_ the time-axis intercept between the tangent at the inflection point and the baseline *Q*_0_. *T*_ini_ was used as a model-derived timing parameter characterizing the early phase of change along the fitted developmental trajectory. Negative *A* values were permitted for decreasing *FD*_rad_ trajectories.

For regional analyses, Gompertz trajectories were fitted separately in each of the 78 CRL cortical regions. Modality-specific observations identified as outliers before fitting were excluded using the original fitting procedure. Regions were retained for the regional timing analysis only when both *FD*_rad_ and curvature trajectories met the original inclusion criteria: a significant overall Gompertz fit and a significant *T*_ini_ parameter after FDR correction, and an estimated *T*_ini_ within the observed gestational-age range. Fifty-two regions met these joint criteria.

Pearson’s correlations were calculated between regional *T*_ini-FDrad_ and *T*_ini-curvature_ values across the 52 retained cortical regions. To account for cortical spatial autocorrelation, statistical significance was assessed using a surface-based spin test with 10,000 rotations. Parcel centroids were derived from the 30-week CRL cortical labels projected onto topology-matched 30-week spherical surfaces. For each rotation, parcel values were reassigned within each hemisphere using a bijective minimum-cost assignment. The same mirrored rotation was applied to the two hemispheres to preserve homologous bilateral organization. A two-sided spin *P* value was calculated as the proportion of rotated correlations with an absolute value at least as large as the observed correlation.

Spatial Pearson’s correlations between *FD*_rad_ and curvature were also calculated across all 78 cortical regions on each weekly atlas from 23 to 38 weeks. Each weekly association was assessed using the same 10,000-rotation surface-based spatial-null procedure. Spin-test *P* values were corrected across the 16 gestational weeks using the Benjamini–Hochberg procedure. As a more stringent family-wise sensitivity analysis, max-|*r*| corrected *P* values were calculated by comparing each observed correlation against the maximum absolute correlation across all 16 weeks in each rotation.

A linear mixed-effects model was fitted to assess the coupled changes in cortical *FD*_rad_ and curvature over the same longitudinal interval:

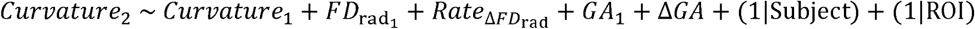

*Curvature*_1, 2_ denote regional curvature measured at the first and second scans, respectively. *FD*_rad1, 2_ denote the corresponding baseline and follow-up *FD*_rad_. *GA*_1, 2_ denote gestational ages at the first and second scans,respectively, and Δ*GA = GA*_2_ – *GA*_1_ denotes the interval between scans. The rate of *FD*_rad_ change was defined as 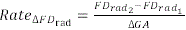 and was expressed per gestational week. All continuous variables were standardized before model fitting. Random intercepts for subject and cortical region were included to account for correlations among measurements from the same fetus and systematic differences among cortical regions. The model included 512 subject–region observations from 16 subjects and 32 cortical regions. Statistical inference was based on Wald z-statistics, with *P* < 0.05 considered statistically significant.

To visualize the overarching within-individual co-developmental patterns, longitudinal *FD*_rad_ and curvature trajectories from all longitudinal scans and cortical regions were aggregated into a 2D state space. This state space, bounded by the global minimum and maximum values of both metrics across the entire dataset, was uniformly partitioned into a 15 × 15 grid of spatial bins. All subjects’ regional developmental displacement vectors within each bin were averaged to compute a consensus vector, color-coded based on trajectory density (i.e., the number of region-subject instances within a given bin) to highlight the dominant developmental pathways (Fig. 5b).

## Data and Code Availability

Source data for all figures, de-identified analysis-level tables, statistical analysis code, and simulation models will be made available at https://github.com/zjuwulab/Fetal-Cortical-Folding. The developing Human Connectome Project (dHCP) data used in this study are publicly available, with access instructions provided at https://biomedia.github.io/dHCP-release-notes/. Raw ZJU-Fetal MRI data, clinical information, and participant-specific images and cortical surfaces cannot be publicly released because of participant privacy and ethical restrictions but may be obtained from the corresponding author upon reasonable request and subject to institutional approval and an appropriate data-use agreement.

## Acknowledgements

This work was supported by the National Natural Science Foundation of China grant nos. 32427802 and U24A20754 to DW, the Key Research Development Project of Zhejiang Province grant no. 2026C02A1262(SD2) to DW, the National Natural Science Foundation of China grant no. 82604061 to RC, and the Postdoctoral Fellowship Program of CPSF grant nos. 2026M793224 and GZC20251895 to RC.

