## Supplementary for "*In Utero* MRI of Radial Glial Fibers Highlights Their Role in Modulating Fetal Cortical Folding"

### Supplementary Material

#### Supplementary Methods

##### Normalized Radiality Analysis

To characterize radial orientation while accounting for the overall apparent fiber-density amplitude, normalized radiality was calculated separately for each diffusion MRI scan and each of the 78 CRL cortical regions as regional *FD*_rad_ divided by the corresponding regional *FD*_total_. *FD*_total_ was derived by summing apparent fiber density across all fixels using the same cortical-plate mask, spatial mapping, and regional parcellation as for *FD*_rad_. The ratio was therefore defined at the regional level rather than calculated independently within each voxel:

$$Normalized {radiality}_{s, r}=\frac{FD_{\mathrm{ra}d_{s,r}}}{FD_{\mathrm{tota}l_{s,r}}}$$

where *s* indexes the diffusion MRI scan and *r* indexes the cortical region. The analysis included 93 diffusion MRI scans from 92 participants, yielding 7,254 scan-region observations. All normalized-radiality values were finite and ranged from 0.377 to 0.902.

Developmental change in whole-cortex normalized radiality was assessed using the same modified four-parameter Gompertz function used for whole-cortex *FD*_rad_: $Q=a+bexp[-exp(-k(GA-T_{\mathrm{ini}})+1)]$, where *a* is the baseline, *b* is the signed trajectory amplitude, *k* is the rate parameter, and *T*_ini_ is the model-derived early transition location. The model was fitted to the whole-cortex mean normalized radiality of each diffusion MRI scan, and parameter uncertainty was derived from the nonlinear least-squares covariance matrix.

The association between normalized radiality and curvature was evaluated in 52 paired sessions from 51 participants with matched diffusion and structural MRI, comprising 4,056 scan-region observations. Curvature, normalized radiality, and gestational age were standardized before model fitting. Curvature was modeled as a function of normalized radiality and gestational age with cortical-region fixed effects, and two-way cluster-robust standard errors were again calculated for participant and cortical region. The normalized-radiality coefficient is therefore reported as a standardized effect adjusted for gestational age and stable regional differences.

##### Longitudinal b = 400 s/mm² analysis

The b = 400 s/mm² sensitivity analysis used the same 16 participants, two sessions per participant, 32 cortical regions, and curvature measurements as the primary longitudinal analysis. Cortical *FD*_rad_ was recalculated from the b = 400 s/mm² data. We fitted the same linear mixed-effects model with crossed random intercepts for participant and cortical region. The primary co-development model included baseline curvature, baseline *FD*_rad_, the per-week rate of *FD*_rad_ change, baseline gestational age, and inter-scan interval. A difference-form sensitivity model replaced the change rate with the unscaled within-subject change in *FD*_rad_. A baseline-only model omitted the *FD*_rad_ change term to test prospective association.

#### Supplementary Results

##### Developmental change in normalized radiality

Whole-cortex normalized radiality followed a decreasing modified Gompertz trajectory, with a model-derived *T*_ini_ of 27.51 ± 0.25 weeks (95% CI [27.02, 28.00 weeks], *R*² = 0.961; Fig. S2a). After adjustment for gestational age and cortical-region fixed effects, normalized radiality remained negatively associated with curvature (standardized *β* = -0.455, two-way cluster-robust SE = 0.0319, 95% CI [-0.518, -0.393], *z* = -14.30, *P* < 0.001; Fig. S2b). Because normalized radiality expresses the radial component relative to total apparent fiber density, these findings provide complementary evidence for a developmental reduction in cortical radial organization and its coupling with cortical curvature.

##### Longitudinal b = 400 s/mm² results

The b = 400 s/mm² analysis reproduced the coupled longitudinal association observed with b = 1000 s/mm². In the change-rate model, the rate of *FD*_rad_ change was negatively associated with follow-up curvature (standardized *β* = -0.203, SE = 0.041, 95% CI [-0.284, -0.122], *z* = -4.92, *P* < 0.001). The difference-form model gave a consistent result for within-subject change in *FD*_rad_ (standardized *β* = -0.280, SE = 0.050, 95% CI [-0.378, -0.181], *z* = -5.57, *P* < 0.001). By contrast, baseline b = 400 s/mm² *FD*_rad_ was not associated with follow-up curvature in the baseline-only model (standardized *β* = 0.003, SE = 0.036, 95% CI [-0.068, 0.073], *z* = 0.07, *P* = 0.942). Thus, the lower-b-value data support coupled within-interval changes but not prospective prediction by baseline *FD*_rad_ (Fig. S2).

#### Supplementary Figures

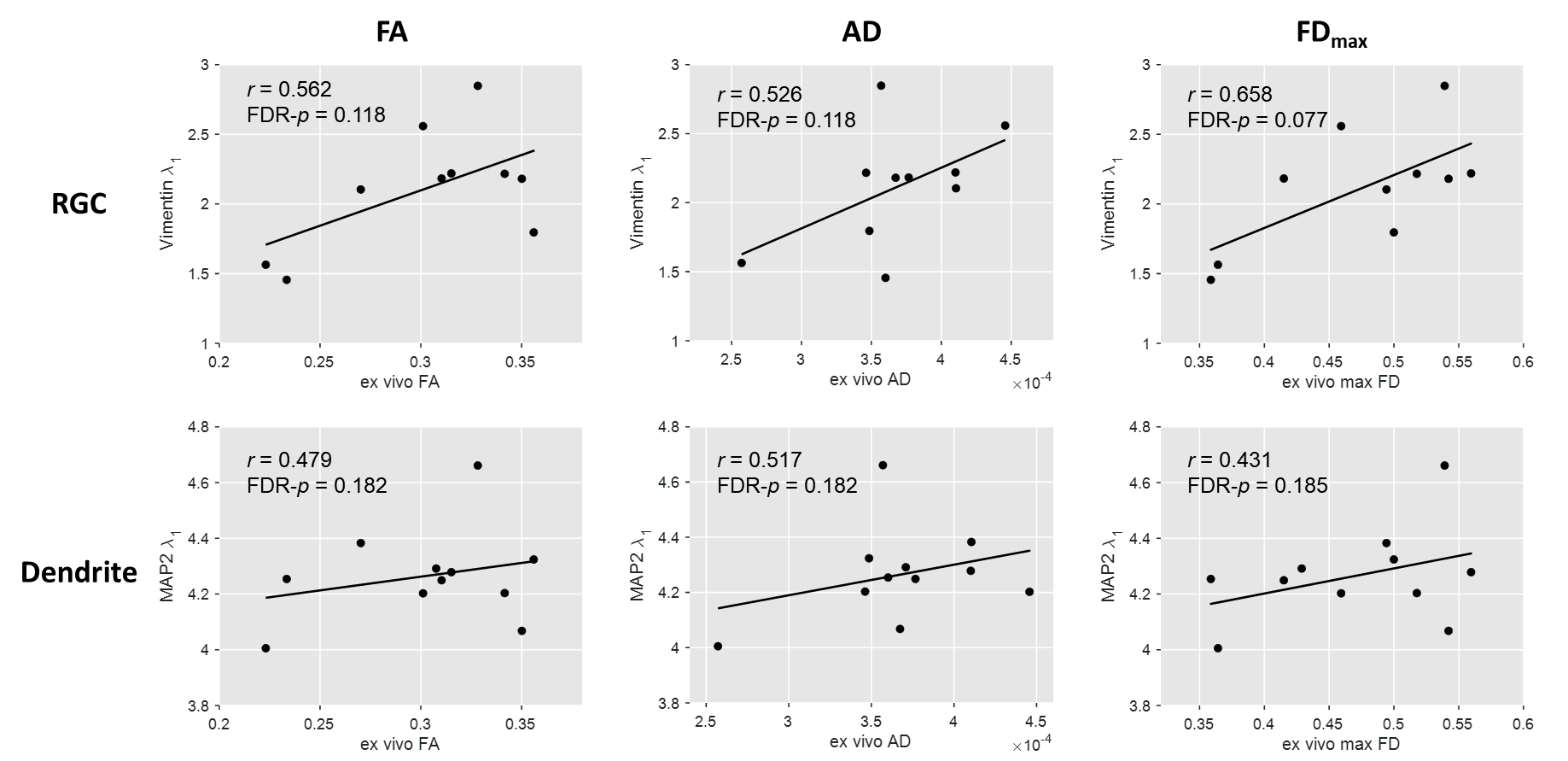

**Fig. S1 | Associations between histology-derived structure-tensor measures and alternative MRI metrics.** No significant correlation was observed in FA, AD, and *FD*_max_ with either vimentin- or MAP2-based structural tensor *λ*_1_ (FDR-corrected P > 0.05).

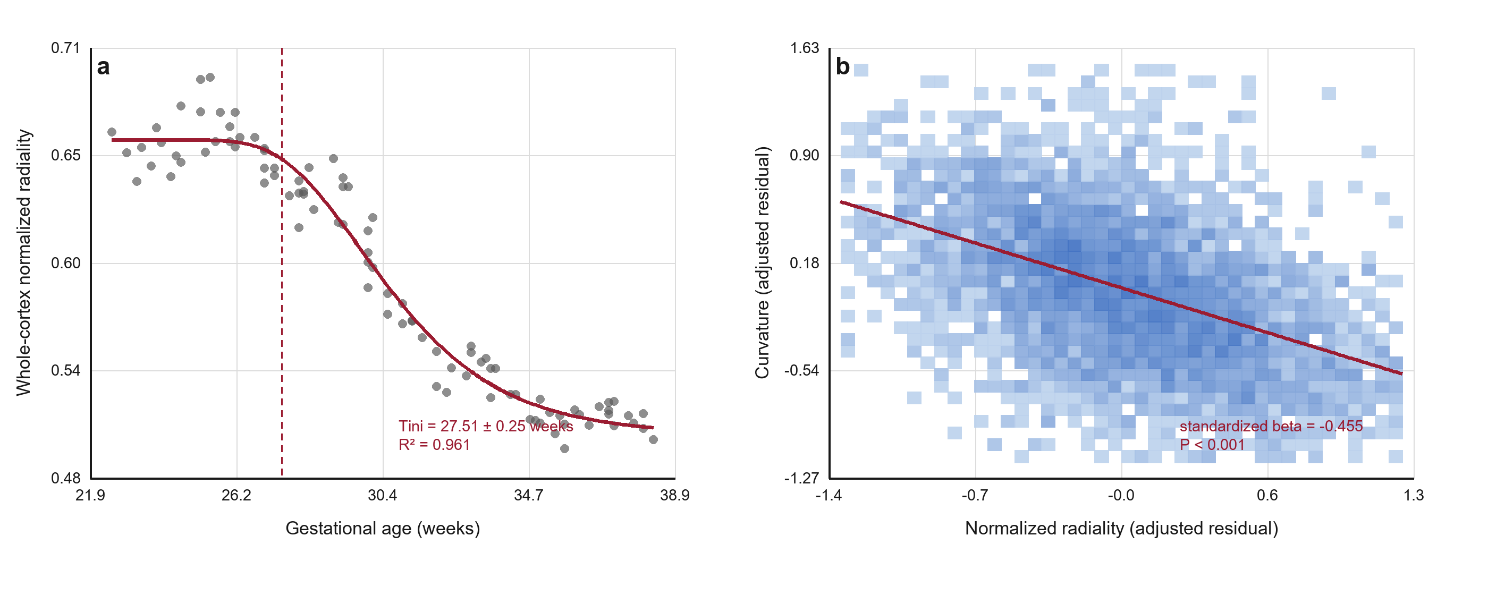

**Fig. S2 | Supplementary normalized radiality analysis. a. Whole-cortex normalized radiality across gestational age.** Each point represents one diffusion MRI scan; the solid line shows the fitted modified Gompertz trajectory, and the dashed vertical line marks its model-derived *T*_ini_. **b. Partial association between normalized radiality and curvature after adjustment for gestational age and cortical-region fixed effects.** The hexagonal bins show observation density, and the line illustrates the fitted negative association. *FD*_total_ was used only as the denominator in the normalized radiality measure; no separate developmental trajectory of *FD*_total_ is reported.

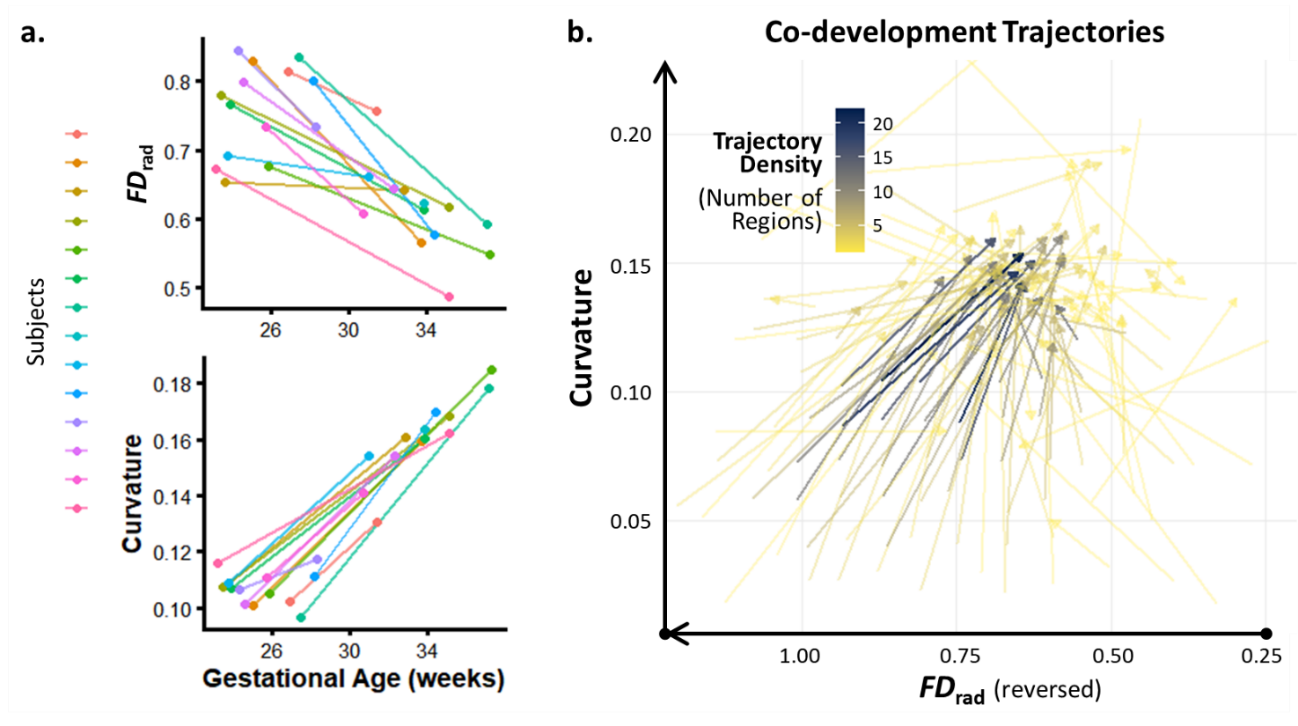

**Fig. S3 | Coupled longitudinal changes in cortical *FD*_rad_ and curvature reproduced using the b = 400 s/mm^2^ data. a. Longitudinal development from mid- to late-gestation.** Longitudinal changes in cortical *FD*_rad_ and curvature were consistent with the cross-sectional observation that decreasing *FD*_rad_ was accompanied by increasing curvature. Only subjects within the studied GA range were displayed (GA > 23 weeks). **b. Co-developmental trajectories of regional *FD*_rad_ and curvature.** Each arrow represents the co-developmental trend of a specific cortical region. The color of the arrows indicates the trajectory density (i.e., the number of regions sharing a similar developmental path), with darker colors representing a higher consensus among regions. The “core development flow” summarized the predominant pattern from a high-*FD*_rad_/low-curvature state and progressed towards a low-*FD*_rad_/high-curvature state.
